# Allelic differences of clustered terpene synthases contribute to correlated intra-specific variation of floral and herbivory-induced volatiles in a wild tobacco

**DOI:** 10.1101/2020.04.26.062133

**Authors:** Shuqing Xu, Christoph Kreitzer, Erica McGale, Nathalie D. Lackus, Han Guo, Tobias G. Köllner, Meredith C. Schuman, Ian T. Baldwin, Wenwu Zhou

**Author notes:** Authors for Correspondence: Shuqing Xu, Institute for Evolution and Biodiversity, University of Münster, Hüfferstrasse 1, Münster, Germany. and Wenwu Zhou, Institute of Insect Sciences, Zhejiang University, Hangzhou, China.

## Abstract

- Plant volatile emissions can recruit predators of herbivores for indirect defence and attract pollinators to aid in pollination. Although volatiles involved in defence and pollinator attraction are primarily emitted from leaves and flowers, respectively, they will co-evolve if their underlying genetic basis is intrinsically linked, either due to pleiotropy or genetic linkage. However, direct evidence of co-evolving defence and floral traits is scarce.
- We characterized intra-specific variation of herbivory-induced plant volatiles (HIPVs), the key components of indirect defence against herbivores, and floral volatiles in the wild tobacco *Nicotiana attenuata*.
- We found that variation of (*E*)-β-ocimene and (*E*)-α-bergamotene contributed to the correlated changes in HIPVs and floral volatiles among *N. attenuata* natural accessions. Intra-specific variations of (*E*)-β-ocimene and (*E*)-α-bergamotene emissions resulted from allelic variation of two genetically co-localized terpene synthase genes, *NaTPS25* and *NaTPS38* respectively.
- Analyzing haplotypes of *NaTPS25* and *NaTPS38* revealed that allelic variations of *NaTPS25* and *NaTPS38* resulted in correlated changes of (*E*)-β-ocimene and (*E*)-α-bergamotene emission in HIPVs and floral volatiles in *N. attenuata*.
- Together, these results provide evidence that pleiotropy and genetic linkage result in correlated changes in defences and floral signals in natural populations, and the evolution of plant volatiles is likely under diffuse selection.

## Introduction

Plant volatile emissions mediate interactions between plants and their friends and foes at distance, which significantly affect plant fitness in nature (Dicke *et al*., 1994; **Baldwin**, 2010). When attacked by herbivores, many plants emit herbivory-induced plant volatiles (HIPVs) from their leaves, which attract predators or parasitoids of herbivores, as well as deter herbivore oviposition, thus reducing herbivore loads and increasing the chance of plant survival (Turlings & Wäckers, 2004; Halitschke *et al*., 2008; Degenhardt *et al*., 2009; Allmann & Baldwin, 2010; Schuman *et al*., 2012). For many flowering plants, floral scent is a key signal that attracts pollinators to transfer pollen, and hence is critical for their reproductive success (Raguso, 2008). Interestingly, many of the same volatile organic compounds are found both in floral volatiles and HIPVs, such as terpenoids (Tholl, 2006; Raguso *et al*., 2016).

Terpenoids represent the largest class of plant-derived natural products (Boutanaev *et al*., 2015), which are produced from terpene precursors: isopentenyl diphosphate, geranyl diphosphate or farnesyl diphosphate, by their “signature enzymes”: terpene synthases (TPSs; Tholl, 2006; Degenhardt *et al*., 2009). Because terpene precursors are commonly found in different plant tissues and species, changes in terpenoid abundance, in both floral volatiles and HIPVs, are largely determined by the expression and function of TPSs (van Schie *et al*., 2006; Tholl, 2006; Nagegowda, 2010; Irmisch *et al*., 2012). Most previous studies investigated only floral and foliar expression of TPSs and terpenoid emissions in isolation. However, Lee et al. found that a single homoterpene synthase, expressed in both *Arabidopsis* flowers and herbivore-induced leaves, resulted in the biosynthesis of (*E,E*)-4,8,12-trimethyltrideca-1,3,7,11-tetraene in both HIPVs and floral volatiles (Lee *et al*., 2010). Furthermore, recent genomic studies showed that many TPSs are co-localized in the genome and form different clusters (defined as two or more genes located near to each other in the genome that were evolved from the same common ancestral gene or involved in a same biosynthetic pathway) (Falara *et al*., 2011; Chen *et al*., 2019; Chen *et al*., 2020), which can result in strong genetic linkages among the different TPSs that are expressed in different tissues. These studies suggest that pleiotropy or genetic linkage can result in co-evolution of floral and foliar terpenoid emissions, which may lead to the correlated changes in plant-pollinator and plant-herbivore interactions. However, direct evidence of this inference remains scarce.

In *Nicotiana attenuata*, a wild tobacco native to the North American Great Basin Desert, the same terpenoids are also found in both HIPVs and floral volatiles (Kessler & Baldwin, 2007; Wu *et al*., 2008; Steppuhn *et al*., 2008; Gaquerel *et al*., 2009; Schuman *et al*., 2009). Analysing the HIPVs in five natural accessions revealed that herbivory-induced (*E*)-β-ocimene and (*E*)-α-bergamotene emissions are variable in *N. attenuata* (Wu *et al*., 2008; Steppuhn *et al*., 2008; Schuman *et al*., 2009). Through manipulation of individual HIPVs in plants, studies showed that (*E*)-α-bergamotene can significantly increase the attraction of predatory bugs to *N. attenuata* (i.e. *Geocoris* spp.) and improve plant fitness under herbivore attack (Kessler & Baldwin, 2001; Halitschke *et al*., 2008; Schuman *et al*., 2012). Moreover, augmented levels of (*E*)-α-bergamotene emission in leaves by ectopic expression of a maize *TPS* gene in *N. attenuata* also suggested that (*E*)-α-bergamotene may interact with green leaf volatiles to improve plant fitness in populations of emitting or non-emitting plants (Schuman *et al*., 2015).

Although *N. attenuata* is a predominantly self-fertilizing species, it opportunistically attracts pollinators for outcrossing (Sime & Baldwin 2003; Kessler *et al*., 2008; Guo *et al*., 2019). The analysis of floral volatiles of *N. attenuata* revealed that both (*E*)-β-ocimene and (*E*)-α-bergamotene are also present (Kessler & Baldwin, 2007). While the emission of (*E*)-α-bergamotene in corolla tubes increases *M. sexta* moth-mediated pollination success (Zhou *et al*., 2017), (*E*)-β-ocimene can elicit distinct electrophysiological responses in the *M. sexta* antennae and may act as a repellent to the moths (Fraser *et al*., 2003; Kessler & Baldwin, 2007). However, whether the emission of (*E*)-β-ocimene and (*E*)-α-bergamotene in HIPVs and floral volatiles show correlated changes among different genotypes remains unknown.

Here, we first screened the floral volatiles and HIPVs of natural accessions collected from different habits within the range of *N. attenuata*’s geographical distribution. We found that (*E*)-β-ocimene and (*E*)-α-bergamotene in HIPVs and floral volatiles showed correlated changes among *N. attenuata* genotypes. Then, using both forward and reverse genetic tools, we characterized the genetic mechanisms underlying the variations of (*E*)-β-ocimene and (*E*)-α-bergamotene in HIPVs and floral volatiles, respectively. The results showed that the within-species variation of (*E*)-β-ocimene and (*E*)-α-bergamotene results from expression changes of two clustered *TPS* genes in the TPS-b clade located on chromosome 2: *NaTPS25* and *NaTPS38*, respectively. To further examine whether the identified two *TPS* genes contributed to the correlated changes of HIPVs and floral volatiles among *N. attenuata* genotypes, we characterized the haplotypes of the *NaTPS25* and *NaTPS38* in *N. attenuata* and their associations with HIPV and floral volatiles. The results demonstrated allelic differences of the *NaTPS25* and *NaTPS38* gene cluster result in correlated changes in both floral volatiles and HIPVs in natural populations of *N. attenuata*.

## Materials and Methods

### Plant materials and growth conditions

Seeds of *Nicotiana attenuata* Torrey ex. Watson (Solanaceae) natural accessions (genotypes) that were collected from a selection of habitats within the range of the species in the southwestern United States (Table S1) and inbred for one generation in the glasshouse in Jena (Germany) except for two inbred lines: Utah and Arizona, which have been self-fertilized for 30 and 22 generations in the glasshouse, respectively. To develop the AI-RIL population, Utah genotype (self-fertilized for 30 generations) and Arizona genotype (self-fertilized for 22 generations) were crossed to generated F1 plants, which were then self-fertilized to generate 150 F2 plants (Zhou *et al*., 2017). Briefly, in each generation from F2 to F6, we intercrossed ∼150 progeny using a random mating and equal contribution crossing design. For generation F7, two seeds from each of the crosses at F6 were germinated and used for the single-seed descendent inbreeding process. In total, five generations of inbreeding were conducted (from F6 to F11).

Seeds were germinated following the protocol described by Krügel *et al*. (Krügel *et al*., 2002): seeds were sterilized, placed on Gamborg’s B5 medium, and kept under LD (16h : 8h, light : dark) in a growth chamber (Percival, Perry, IA, USA) for 10 d, and then transferred to small TEKU pots for another 10 d in the glasshouse. Subsequently, plants were transferred to 1 l pots and kept in the glasshouse (26 ± 1°C; 16h : 8h light : dark). For the VIGS experiments, plants in 1 l pots were transferred to a climate chamber under a constant temperature of 26 °C, a 16h : 8h (light : dark) light regime and 65% relative humidity (Galis *et al*., 2013). Watering, fertilization, and light regimes were previously described (Galis *et al*., 2013; Schuman *et al*., 2014).

### Measuring leaf volatiles among 25 natural genotypes and AI-RILs

To measure the emission of leaf volatiles from the 25 natural genotypes, plants in the elongation stage (50 d after germination) were used. Simulated herbivory treatments and leaf volatiles collection were conducted as described in detail in a previous study (Zhou *et al*., 2017). Briefly, the simulated herbivory treatment was carried out at 8 am, the leaves were wounded with a pattern wheel, and 20 µl diluted (1:5 dilution with water) *M. sexta* oral secretion was supplied to the wounds. To obtain temporally resolved comparative data for leaf volatiles, small pieces of polydimethylsiloxane tubes (see Kallenbach *et al*., 2014) were incubated with the damaged leaves from 12-4 pm, 4-8 pm and 8pm-6 am (to the following morning). To measure the effects of jasmonic acid signalling on the induction of leaf volatiles in the 25 genotypes, plants in the elongation stage (55 d after germination) were treated with MeJA. At 8 am, 150 µg MeJA dissolved in 40 µl lanolin paste was deposited at the base of one stem leaf of each plant. After 24 h of the MeJA treatment, the leaf was enclosed into a plastic cup (polyethylene terephthalate) together with two polydimethylsiloxane tubes, and the polydimethylsiloxane tubes were collected 24 h later and kept in -20 °C until analysis. Three biological plant replicates were sampled for each group.

The measurement of leaf volatiles from the AI-RIL plants (55 d after germination) has been previously described (Zhou *et al*., 2017). Briefly, simulated herbivory treatment was carried out at 8 am, and the leaf was immediately enclosed in to a plastic cup where two polydimethylsiloxane tubes were incubated as described above. The polydimethylsiloxane tubes were collected after 24 h (8 am the next day), and immediately kept at -20 °C until analysis.

### QTL mapping

To identify genetic basis underlying HIPV variations in *N. attenuata*, we deployed a forward genetic mapping approach. The QTL mapping was described in detail in our previous study (Zhou *et al*., 2017) and was conducted with the R package QTLRel (Cheng *et al*., 2011) following the tutorial. In brief, the genotype information of AI-RIL plants and the linkage map were obtained from a dataset reported earlier (Guo *et al*., 2020, Zhou *et al*., 2017). In total, we obtained 25,469 SNP markers among 248 individuals. These markers were grouped into 12 linkage groups. The peak area of herbivore-induced (*E*)-β-ocimene was log-transformed. Samples with missing genotype or phenotype information were removed. The variance of (*E*)-β-ocimene within the population was estimated via ‘estVC’, and then used for the genome-wide scan together with the genotypes. The empirical threshold was estimated based on 500 permutations.

### Sequencing and assembling the leaf transcriptome of the Arizona genotype

To identify candidate genes that are associated with (*E*)-β-ocimene biosynthesis, we sequenced the transcriptomes of both control and *M. sexta* oral secretion-induced rosette leaves using Illumina HiSeq 2000. The growth conditions and simulated herbivory treatments were as described above. Samples from untreated plants were used as control. All samples were collected at noon. Total RNA was extracted from ∼100 mg of plant tissue using TRIzol (Thermo Fisher Scientific) according to the manufacturer’s protocol. All RNA samples were subsequently treated with DNase-I (Fermentas) to remove all genomic DNA contamination. The mRNA was enriched using an mRNA-seq sample preparation kit (Illumina), and ∼ 200 bp insertion size libraries were constructed using the Illumina whole transcriptome analysis kit following the manufacturer’s standard protocol (Illumina, HiSeq system). All libraries were then sequenced on the Illumina HiSeq 2000 at the sequencing core facility at Max Planck Institute for Molecular Genetics (Berlin). After removing the adaptor sequences and low-quality reads, in total, we obtained 25297814 and 19124690 paired-end 100 bp reads for control and induced samples, respectively. The raw reads are deposited in the NCBI SRA repository (SRX1804553 and SRX1804540).

The RNA-seq reads from control and induced samples were pooled for the *de novo* assembly of the leaf transcriptome from Arizona genotype using the trinity tool kit (Grabherr *et al*., 2011). The minimum contig length was set at 200 bp. In total, we assembled 62132 transcripts with N50 length equal to 1364 bp.

### Measuring abundance of *NaTPS25* transcripts in natural accessions

To measure the abundance of *NaTPS25* transcripts in the 25 natural accessions, plants in the elongation stage (50 d after germination) were used. Simulated herbivory treatment was carried out on one stem leaf of each plant at 8 am. Eight hours after treatment, the leaf was harvested and immediately kept in -80 °C until use. Total RNA was isolated from the leaves using TRIzol reagent (Thermo Fisher Scientific) and then treated with DNase-I (Fermentas) following the manufacturers’ instructions, to remove contamination from genomic DNA. One µg total RNA from each sample was reverse transcribed into cDNA using SuperScript II reverse transcriptase following the manufacturer’s instructions (Thermo Fisher Scientific). The relative transcript accumulation of *NaTPS25* was measured using RT-PCR on a Stratagene MX3005P PCR cycler (Stratagene). The elongation factor-1A gene, NaEF1a (accession number D63396), was used as the internal standard for normalization (Oh *et al*., 2013). PCR amplification was performed with Phusion Green High-Fidelity DNA polymerase (Thermo Fisher Scientific) using 40 PCR cycles to enable the detection of rare transcripts of *NaTPS25*. The primers used are given in Table S2.

### Silencing *NaTPS25* using virus-induced gene silencing (VIGS), and measurement of HIPVs from VIGS plants

To investigate the function of *NaTPS25 in vivo*, VIGS based on the tobacco rattle virus was used to transiently knock down the expression of the *NaTPS25* gene in *N. attenuata* as previously described (Galis *et al*., 2013). A 163 bp fragment of the open reading frame region of *NaTPS25* was amplified by PCR with the primers listed in Table S2. PCR fragments were separated by agarose gel electrophoresis, excised and purified by a gel band purification kit (Amersham Biosciences) according to the manufacturer’s instructions. Subsequently, they were digested with BamH1 and Sal1 and inserted into plasmid pTV00. After sequencing, the VIGS constructs and pTV00 (empty vector, EV) were separately transformed into *Agrobacterium*. Additionally, the pTVPDS vector, harbouring a part of the sequence of a phytoene desaturase, was used to monitor the progress of the gene silencing.

The VIGS experiments were carried out on the Arizona genotype. The inoculation was performed 25 d after germination, and the volatile trapping another 20 d later, after the establishment of the viral vector. Before any treatment, three leaves (1/plant) were sampled to test silencing efficiency. Afterwards another three leaves (1/plant) were treated with 75 µg methyl jasmonate dissolved in 20 µl lanolin paste on the base of leaves at 12 pm. After 1 hour, the outer portion of these leaves were harvested and frozen to test the silencing efficiency, after which the whole plant was again trapped for volatile collection. Lanolin paste (20 µl) was used for the control group.

### Cloning and heterologous expression of NaTPS25

To examine the enzymatic function of *NaTPS25 in vitro*, a truncated version of *NaTPS25_AZ* (missing the first 13 codons) was cloned from cDNA of Arizona genotype plants using the primer pair TPS25f/r (Table S2). The resulting PCR fragment was cloned into the pET200/D-TOPO vector (Invitrogen, Carlsbad, CA, USA) and the *Escherichia coli* strain BL21 Codon Plus (Invitrogen) was used for heterologous expression. The protein expression and bacterial extract preparation were done as described in a previous study (Zhou *et al*., 2017). Enzyme assays were performed in a Teflon^®^-sealed, screw-capped 1 ml GC glass vial containing 50 µl of the bacterial extract and 50 µl assay buffer with 10 µM geranyl diphosphate, 10 mM MgCl_2_, 0.2 mM Na_2_WO_4_ and 0.1 mM NaF. A solid phase micro-extraction (SPME) fiber coated with a 100 µm-thick layer of polydimethylsiloxane (SUPELCO, Belafonte, PA, USA) was placed into the headspace of the vial where it was incubated for 60 min at 30°C, before it was inserted into the injector of the gas chromatographer to analyze the absorbed reaction products. GC-MS analysis was conducted using an Agilent 6890 Series gas chromatographer coupled to an Agilent 5973 quadrupole mass selective detector. The GC was operated with a DB-5MS column (Agilent, Santa Clara, USA, 30 m x 0.25 mm x 0.25 µm).

### Targeted resequencing of the cDNA and genomic regions of *NaTPS38* and confirmation of the inversion of *NaTPS25* in Utah genotype

To sequence the *NaTPS38* locus from all the accessions and characterize its allelic variations, we amplified the full-length open reading frame using the high-fidelity Phusion DNA Polymerase (Thermo Fisher Scientific) and the primer combination TPS38f and TPS38r (Table S1). The following cycle parameters were used: 98°C for 30 sec; 35 cycles with 98°C for 10 s, 52°C for 20 s, 72°C for 1.5 min and a final step with 72°C for 5 min. PCR products were separated by agarose gel electrophoresis, excised and purified with a PCR clean-up gel extraction kit (Macherey-Nagel), then cloned into the pJET 1.2/blunt vector (Thermo Fisher Scientific) and transformed into *E. coli* (TOP10 Competent Cells, Invitrogen) according to the manufacturers’ instructions. Plasmid extraction was performed using the NucleoSpin Plasmid Kit (Macherey-Nagel). Sequencing reactions were performed using the BigDye Terminator mix (Thermo Fisher Scientific). The obtained sequences were manually corrected, and data analysis was executed with Geneious^®^6.0.5. The cDNA sequences are deposited in the NCBI database (accession numbers: MN400683 -MN400708).

To examine the genetic architecture of *NaTPS25*, we isolated genomic DNA from 10-d-old seedlings using a cetrimonium bromide (CTAB) extraction buffer (Bubner *et al*., 2004). Genomic DNA purity and integrity were evaluated by amplifying the elongation factor-1A with the primer set shown in Table S2. Sequencing approaches for the comparison between Arizona genotype and traced fragments in Utah genotype were conducted as described above. The PCR fragment amplifications were carried out using the Phusion Green DNA Polymerase (Thermo Fisher Scientific) in 20 µl PCR reactions (1 x Phusion Green HF Buffer, 10 mM dNTPs, 3% DMSO and 5 mM primers). The PCR program (30 s at 98°C, 35 cycles of 10 s at 98°C, 30 s at 60°C, 2 min at 72°C with a final elongation of 5 min at 72°C) varied in the annealing temperature (72°C) according to the primer set (Table S2). We validated the presence of the intact *NaTPS25* allele with primer set I (PrS I), and the presence of exon 6-7 with primer set II (PrS II). Moreover, we utilized two primer combinations to provide genetic information for the inversion event of exon 6 and 7. Primer set II (PrS II) was used to detect exons 6 and 7, and primer set III (PrS III) to amplify a fragment specific to this inversion.

### Measuring the abundance of *NaTPS38* transcripts in hybrid plants

To characterize whether the variations of transcript abundance of *NaTPS38* were due to changes in *cis*-or *trans*-regulatory elements, we measured the abundance of *NaTPS38* transcripts in the two parental inbred lines and their hybrid lines in F1. Plants in their elongation stage (50 d after germination) were used. Simulated herbivory treatment was carried out at 8 am on one stem leaf of each plant. Eight hours after treatment, leaves were harvested and flash-frozen in liquid nitrogen, then kept at -80 °C until processing. Total RNA isolation, cDNA synthesis and qPCR amplification were carried out as described above. The primers used to specifically amplify the alleles are shown in the supplementary file (Table S2).

### Statistical analysis

All statistical analyses were performed in R 3.5.1 (R Core Team, 2016). The functions of “prcomp”, “t.test” and “anova” from the “stats” package were used for principal components analysis, student’s *t*-tests and ANOVA analysis, respectively. The raw data and data analysis R scripts are deposited on figshare repository: https://figshare.com/s/56b11cee270eb04f7ecb.

## Results

### (*E*)-β-ocimene and (*E*)-α-bergamotene contributed to associated changes in HIPVs and floral volatiles among *N. attenuata* natural accessions

We first investigated HIPVs variations in *N. attenuata* by sampling the herbivory-induced leaf headspace from 25 genotypes that were collected from different habitats within the species range. We found two monoterpenes, (*E*)-β-ocimene and linalool, and one sesquiterpene, (*E*)-α-bergamotene, to be highly variably released among the studied genotypes (Fig. 1**a-c**). In addition, previous studies showed that the emission of all three compounds in leaves were significantly induced by herbivory (Halitschke *et al*., 2000), supporting their classification as HIPVs. Jasmonate signalling is involved in regulating the emission of many HIPVs in *N. attenuata*, and its accumulation is variable among wild genotypes (Schuman *et al*., 2009). To examine whether the observed HIPVs variations were attributable to differences in herbivory-induced jasmonate signalling, we supplied a similar level of methyl jasmonate to the leaves of these genotypes. Under similarly elevated levels of jasmonate signalling in methyl jasmonate-treated plants, the emission of the three terpenoids remained variable and were the major contributors to the variation in HIPVs among the investigated genotypes (Fig. 1**d**). This suggests that intra-specific variations of HIPV is largely independent of jasmonate signalling.

**Fig. 1.**
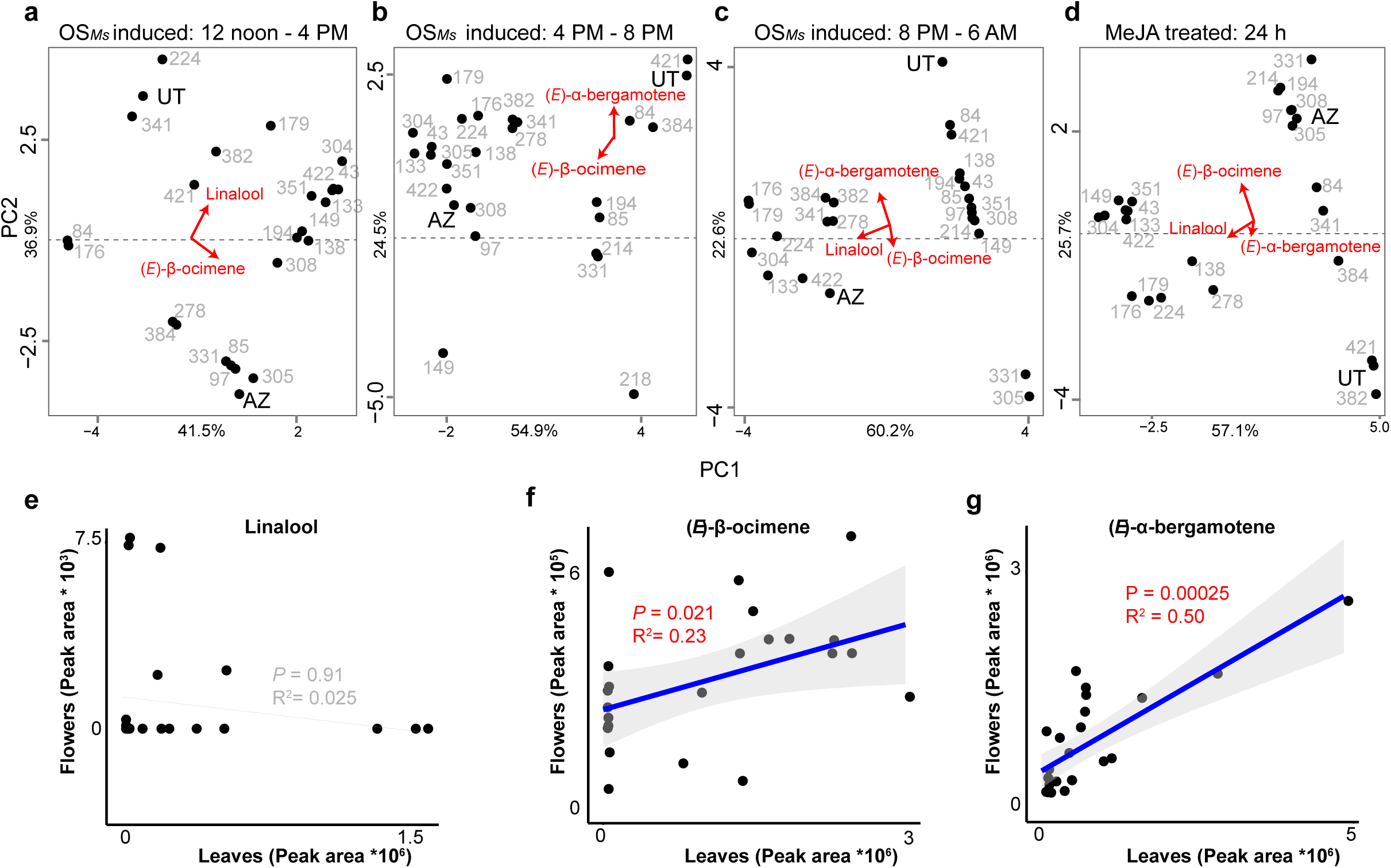
Variations of HIPVs and correlated changes between HIPVs and floral volatiles. (**a-d**) Principle component analysis of leaf volatiles after simulated herbivory (a-c), or after MeJA treatment (d). For simulated herbivory treatment (OS*MS*), leaves were treated with wounding and oral secretion of *M. sexta* at 8 am, and headspace compositions were sampled over three different time periods (12-4 pm, 48 pm and 8pm - 6am) after the OS induction. For MeJA treatment, volatiles were sampled from leaves in a single incubation, 24-48 h after treatment. Variance explained by PC1 and PC2 is shown on x-axis and y-axis, respectively. The four plots should not be compared among each since the variance / covariance structure of the data is not the same. (**e-g**) correlated emission changes between flowers and methyl jasmonate treated leaf headspaces among different genotypes were found in (*E*)-α-bergamotene and (*E*)-β-ocimene but not linalool. Each dot represents a genotype. Spearman rank correlations were used for estimating *P*-values and R^2^. Ribbons refer 95% confidence intervals.

In addition to a flower-specific volatile, benzoyl acetone (Guo *et al*., 2020), the three terpenoids in HIPVs, (*E*)-β-ocimene, linalool, and (*E*)-α-bergamotene were also found in *N. attenuata* floral volatiles (Kessler *et al*., 2007). We reasoned that if the same compound in HIPVs and floral volatiles was regulated by the same genetic mechanism, its abundance would show correlated changes in HIPVs and floral volatiles amounts among different genotypes. To examine this, we further measured the floral volatiles of the 25 genotypes and compared the emissions of (*E*)-β-ocimene, linalool and (*E*)-α-bergamotene emission in the HIPVs and floral volatiles. We found that linalool, which was found in both floral volatiles and HIPVs, showed no correlation among the different genotypes, consistent with a recent study (He *et al*., 2019). However, the variations of (*E*)-α-bergamotene and (*E*)-β-ocimene emission in HIPVs and floral volatiles were significantly correlated (Figure 1; (*E*)-β-ocimene: R^2^ = 0.23, *P* = 0.021; (*E*)-α-bergamotene: R^2^ = 0.50, *P* < 0.001).

### Changes in *NaTPS25* and *NaTPS38* expression are associated with intra-specific variations of (*E*)-β-ocimene and (*E*)-α-bergamotene emissions, respectively, in both HIPVs and floral volatiles

To characterize the genetic mechanisms underlying the correlated changes of (*E*)-α-bergamotene and (*E*)-β-ocimene emission in HIPVs and floral volatiles, we used both forward and reverse genetic tools.

Previously we established an advanced inter-cross recombinant inbred line (AI-RIL) population that was derived from crossing the Arizona and Utah genotypes whose (*E*)-β-ocimene and (*E*)-α-bergamotene emissions differed in HIPVs and floral volatiles. Using this population, we identified a terpene synthase, *NaTPS38*, on linkage group 2 that controls the emissions of (*E*)-α-bergamotene in both HIPVs and floral volatiles (Zhou *et al*., 2017). Among different *N. attenuata* genotypes, the changes in *NaTPS38* expression were correlated with (*E*)-α-bergamotene emissions in both HIPVs and floral volatiles (Zhou *et al*., 2017).

Here, using a similar approach, we found that variations in (*E*)-β-ocimene emission in HIPVs were mapped to the same locus as *NaTPS38* on linkage group 2 of *N. attenuata* (Fig. 2**a** and Zhou *et al*., 2017). This location overlaps with a terpene synthase cluster, which contains a subclade of TPS-b monoterpene synthase genes, including *TPS25, TPS27* and *TPS38* (Falara *et al*., 2011). Among these genes, *TPS25* and *TPS27* in tomato were highly similar to the grape *VvTPS47* that produces (*E*)-β-ocimene. However, after searching the draft genome of *N. attenuata* (created using the Utah genotype), we did not find any genes that showed high similarity to either *TPS25* or *TPS27*. Sequencing the transcriptome of the Arizona genotype allowed us to identify *NaTPS25*, which showed high similarity to *TPS25* and *TPS27* in tomato.

**Fig. 2.**
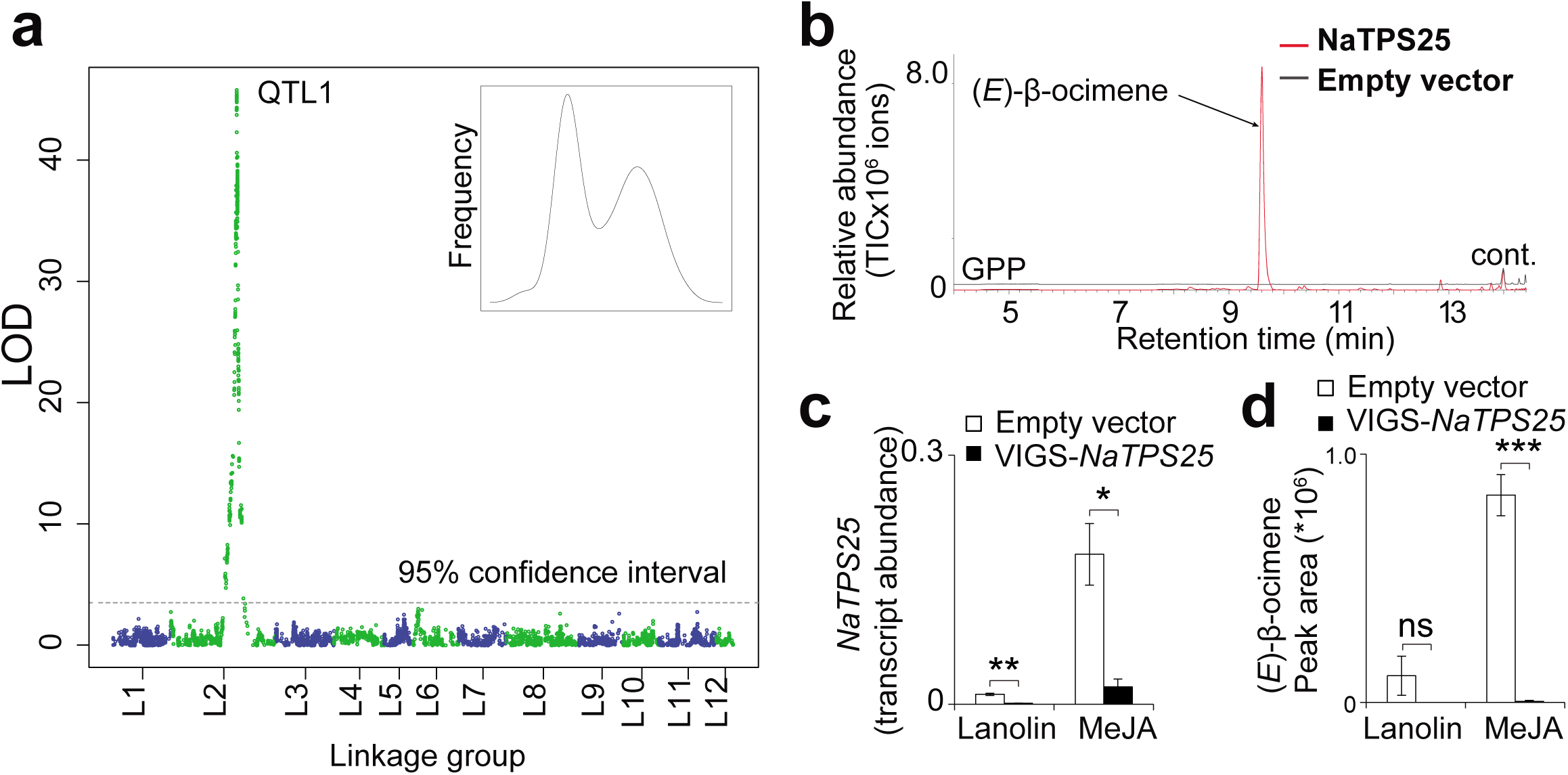
QTL mapping for (*E*)-β-ocimene and functional validation of *NaTPS25*. (**a**) One QTL locus on linkage group 1 (QTL1) for (*E*)-β-ocimene. The 95% confidence interval is marked with a dashed line. (**b**) Heterologously expressed NaTPS25 protein produces (*E*)-β-ocimene *in vitro*. Total ion chromatograms (GC-MS) of the empty vector control (black solid line) and the enzyme products of NaTPS25 protein (red solid line). The enzyme was expressed in *E. coli* and incubated with the substrate geranyl diphosphate. (**c-d**) VIGS-mediated silencing of NaTPS25 expression in leaves. Both constitutive (Lanolin) and methyl jasmonate-induced levels of *NaTPS25* transcript abundance (c) and (*E*)-β-ocimene emission (d) were measured (n=4). Mean and SE are shown. Statistical differences are determined by Student’s *t*-test. *: *P* < 0.05; **: *P* < 0.01; ***: *P* <0.001.

To examine the enzymatic function of *NaTPS25*, we measured the TPS activity of *NaTPS25 in vitro* by heterologous expression of the gene in *Escherichia coli*. The recombinant NaTPS25 protein converts the substrate geranyl diphosphate into (*E*)-β-ocimene as its only product (Fig. 2**b**). To examine the function of *NaTPS25 in vivo*, we silenced the expression of *NaTPS25* in the *N. attenuata* (Arizona genotype) using virus-induced gene silencing (VIGS). When *NaTPS25* expression was silenced, levels of both constitutive and methyl jasmonate induced (*E*)-β-ocimene emission in HIPVs were strongly reduced (Fig. 2**c-d**). Together, these results suggest that herbivore-induced (*E*)-β-ocimene is associated with the expression of *NaTPS25* in HIPVs.

We further investigated whether the expression changes in *NaTPS25* are associated with (*E*)-β-ocimene in HIPVs and floral volatiles. We first measured the presence of *NaTPS25* transcripts among the 25 genotypes by reverse-transcription (RT)-PCR. Interestingly, we observed presence and absence variations of *NaTPS25* at the level of transcripts, which correlate with the emission of (*E*)-β-ocimene from induced leaves among genotypes (Fig. 3**a**). While *NaTPS25* transcripts can be found in all 15 genotypes that emitted (*E*)-β-ocimene after simulated herbivory, they were not found in any of the other 10 genotypes that did not emit detectable levels of (*E*)-β-ocimene (Fig. 3**a**). We then used quantitative polymerase chain reactions (qPCR) to measure the transcript abundance of *NaTPS25* in flowers in order to compare them to emissions of (*E*)-β-ocimene in the respective floral volatile profiles of each genotype. We found that among different genotypes, emissions of (*E*)-β-ocimene in floral volatiles is significantly correlated with *NaTPS25* transcript abundance (*P* = 0.013, Fig. 3**b**). Interestingly, while most of the genotypes that had a high abundance of *NaTPS25* transcripts also had a high floral (*E*)-β-ocimene emission, a few genotypes (e.g., genotype Utah, 43 and 351) showed very low levels of *NaTPS25* transcript abundance, yet still emitted high levels of (*E*)-β-ocimene in their flowers (Fig. 3**b**). These results suggest that in addition to *NaTPS25*, other genes are involved in floral (*E*)-β-ocimene emission. Consistently, in the *N. attenuata* genome (Utah genotype), there is another member of the TPS-b subfamily, of which several homologues from *Vitis vinifera* and *A. thaliana* can synthesize (*E*)-β-ocimene *in vitro* using geranyl diphosphate as the substrate (Huang *et al*., 2010; Martin *et al*., 2010). Furthermore, one homologue of *TPS19* (NIATv7_g29416), which is a member of TPS-e/f subfamily that can produce (*E*)-β-ocimene using NPP as its substrate (Matsuba *et al*., 2013), was found highly expressed in the flowers of *N. attenuata* (http://nadh.ice.mpg.de/NaDH-eFP/cgi-bin/efpWeb.cgi; Brockmöller *et al*., 2017).

**Fig. 3.**
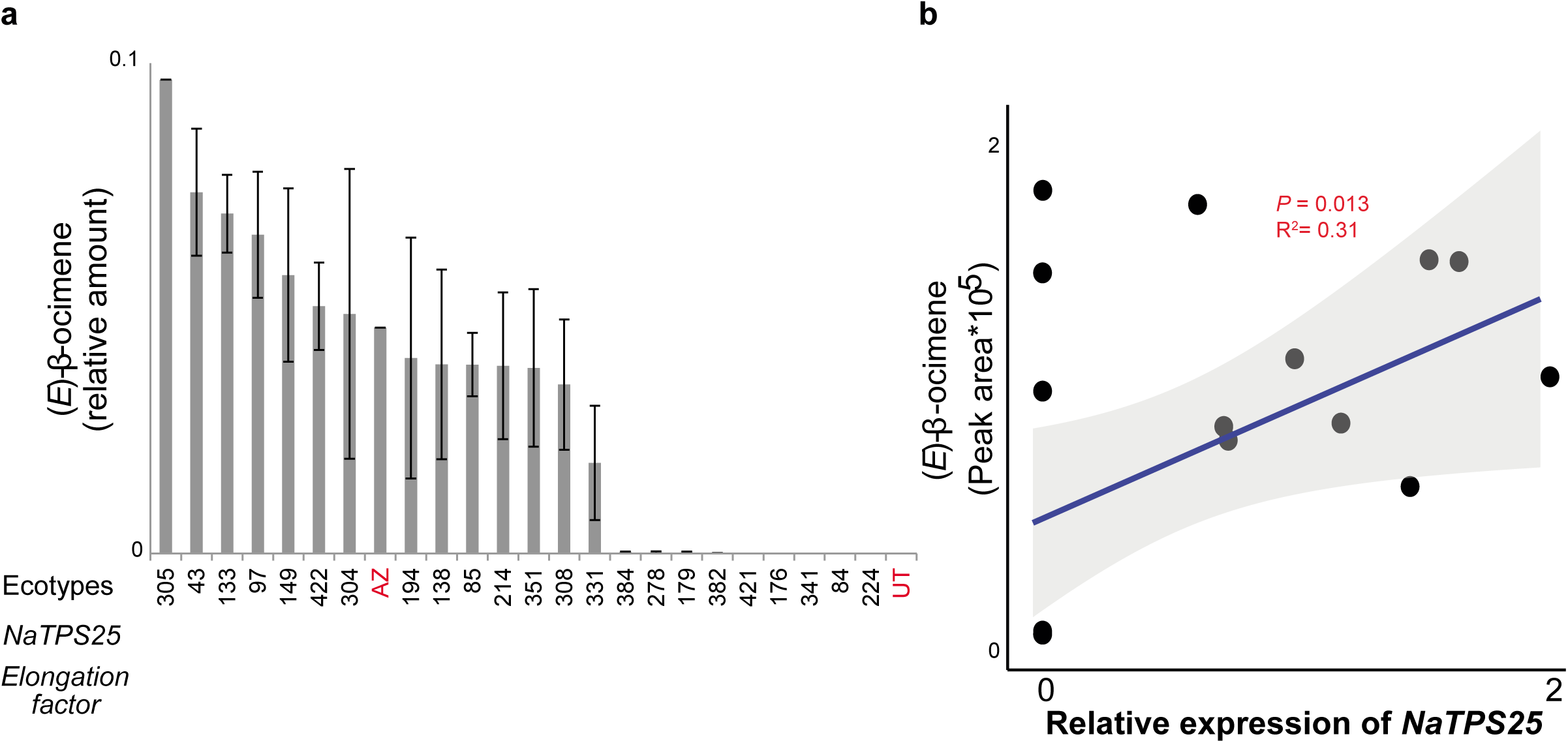
Correlation between transcript abundance and (*E*)-β-ocimene emission. (**a**) Expression of *NaTPS25* and (*E*)-β-ocimene emission in herbivory-induced leaves. X-axis: the names of accessions and the RT-PCR result of *NaTPS25* and *elongation factor-1A* (housekeeping gene); y-axis: the relative amount of (*E*)-β-ocimene in the HIPVs (normalized to all compounds). The leaves were wounded and treated with oral secretion of *M. sexta* larvae at 8 am, volatiles were collected from 12-4 pm (n=3) and leaf samples for expression analysis (RT-PCR for *NaTPS25*) were collected at 4 pm on the same day (n=3). Mean and SE are shown. (**b**) Expression of *NaTPS25* and (*E*)-β-ocimene emission in flowers. Spearman rank correlations were used for estimating *P*-values and R^2^. Ribbons refer 95% confidence intervals.

Together with our previous study (Zhou *et al*., 2017), these results suggest that expression changes in *NaTPS25* and *NaTPS38* are associated with intra-specific variations of (*E*)-β-ocimene and (*E*)-α-bergamotene emissions, respectively, in HIPVs and floral volatiles.

### Two distinct genetic mechanisms resulted in changes in allelic expression of *NaTPS25* and *NaTPS38*

To investigate the genetic mechanisms underlying the allelic expression differences in *NaTPS25* and *NaTPS38*, we analysed sequence polymorphisms of *NaTPS25* and *NaTPS38* among different genotypes.

Consistent with its presence and absence of detectable transcripts levels, we found a structural variation in the *NaTPS25* gene among different genotypes. For genotypes that emit (*E*)-β-ocimene and express *NaTPS25* after simulated herbivory or methyl jasmonate treatment, the complete *NaTPS25* gene can be amplified from genomic DNA with primers that bind to regions containing the start codon and stop codon (Fig. 4**a** & b). There was no obvious gene size variation among these genotypes (Fig. 4**b**). However, for the accessions that failed to emit (*E*)-β-ocimene, *NaTPS25* cannot be amplified using the same primers. Designing exon-specific primers and comparing the genomic sequences of *NaTPS25* revealed that the Utah genotype contains two fragments of *NaTPS25*, disrupted by a ∼9kb long-terminal repeat (LTR)-gypsy transposon. The first fragment contains only exon 6 and 7, whereas the second fragment contains partial sequences of exon 2 and exon 7 (Fig. 4**a**). Using two primer pairs that can specifically amplify these two fragments among different genotypes, we found that most of the genotypes that failed to emit (*E*)-β-ocimene after simulated herbivory, shared the same disrupted gene structure (Fig. 4**c**). Furthermore, additional genomic changes to the *NaTPS25* fragments were found in some genotypes, such as genotype 421, in which the joined fragments between exon 2 and exon 7 could not be amplified, and genotype 176, 341, and 384, in which the exon 6 and 7 fragments could not be amplified. These results showed that allelic differences of *NaTPS25*, caused by LTR-gypsy transposon insertions, contributed to the variations of (*E*)-β-ocimene emission in the HIPVs among different genotypes.

**Fig. 4.**
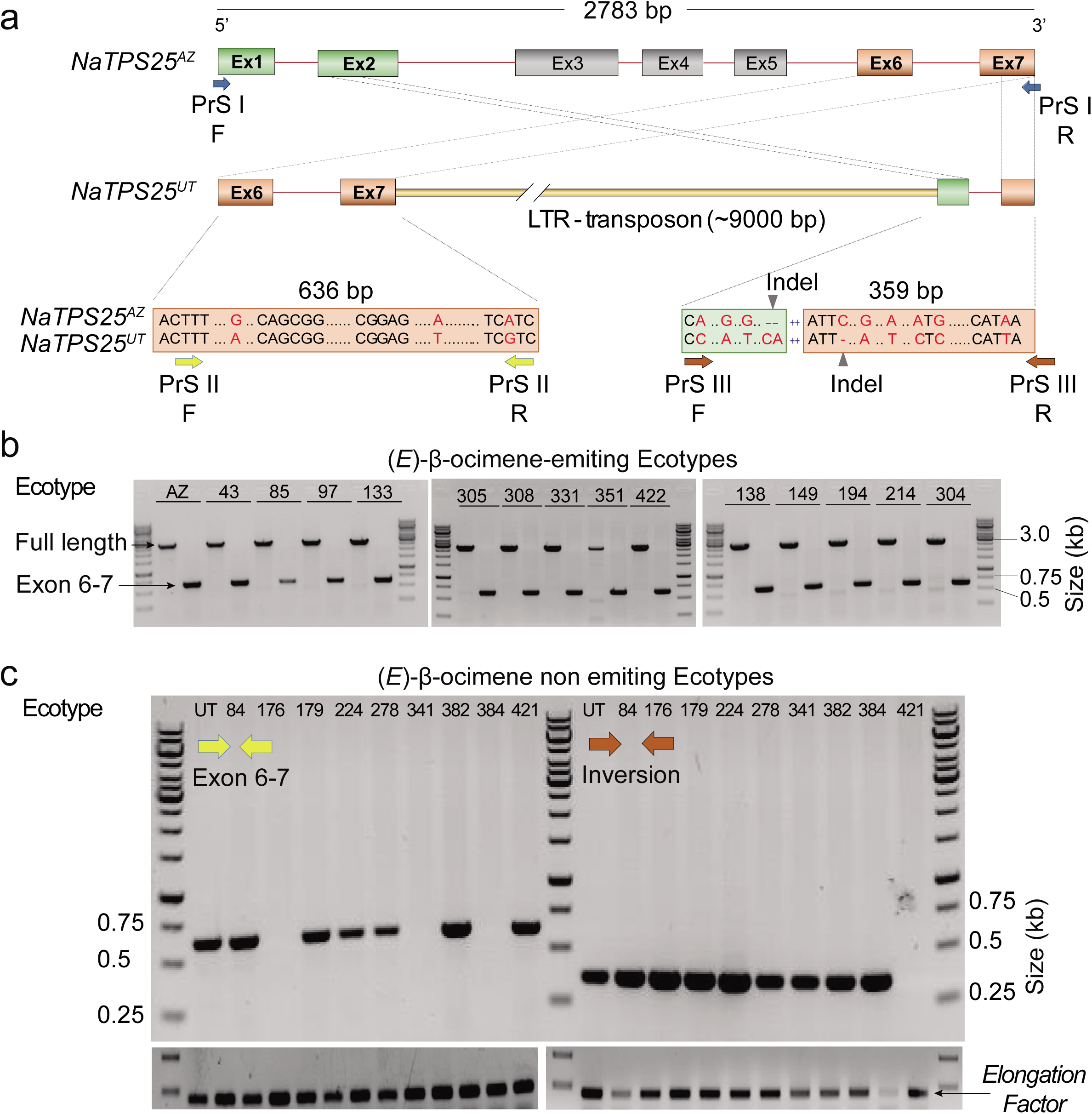
Allelic variation of *NaTPS25* among *N. attenuata* natural accessions. (**a**) The structure of the complete and the inverted *NaTPS25* alleles in *N. attenuata*. The active *NaTPS25* allele in Arizona genotype consists of seven exons (rectangles, Ex1-7) and six introns (red lines). A disruption in the *NaTPS25* allele of the Utah genotype was found by an insertion of a ∼9 kb LTR-gypsy transposon (discontinuous yellow rectangle). Dashed arrows illustrate partially inverted fragments in Utah genotype, in comparison to the allele in Arizona genotype. Alignment of detected fragments in Utah and Arizona genotypes suggested four SNPs (red letters). Two descended regions of the active allele (parts of exon 2 and 7) are linked with a region whose origin is unclear and forms the second fragment (green and orange boxes). Dots refer to the consensus sequences. (**b**) The full length *NaTPS25* PCR fragment was amplified in (*E*)-β-ocimene-emitting genotypes (HIPVs). Primerset (PrS) I (blue arrows) and II (yellow green arrows) were used to amplify the full length and exon 6-7, respectively. (**c**) Presence/absence of exon 6-7 and the inversion region among non-(*E*)-β-ocimene-emitting genotypes (HIPVs). Fragments of exon 6-7 and the inversion region were amplified with primer sets II (yellow green arrows) and III (brown arrows), respectively.

*NaTPS38* transcripts were found in all genotypes but differed in their abundances (Zhou *et al*., 2017). To test whether polymorphic sites at a *cis*-regulatory region and/or changes in *trans*-regulatory elements contributed to the observed variations in the expression of *NaTPS38*, we measured the abundance of *NaTPS38* transcripts in the plants from Arizona and Utah genotypes and their F1 progeny. Because all of the *trans* factors are shared amongst the F1 progeny, differences in allelic transcript abundance between parents and F1 indicate changes in relative *trans*-regulatory activity, whereas differences in relative transcript abundance between two alleles in F1 indicate changes in relative *cis*-regulatory activity (Wittkopp, 2013). After simulated herbivore elicitation, while transcript levels of the *NaTPS38* Arizona allele did not differ between parental and F1 plants (Fig. 5, *P* > 0.1), the levels of the *NaTPS38* Utah allele were significantly lower in F1 plants than in Utah genotype (*P* = 0.0079, Student’s *t*-test), suggesting that the Arizona genotype carry a *trans*-suppressor of *NaTPS38*. In addition, the relative transcript abundances of Arizona and Utah alleles in F1 plants were significantly different (Fig. 5, paired Student’s *t*-test, *P* = 0.012), indicating that the *cis*-regulatory activity of *NaTPS38* is different between Arizona and Utah genotypes. We further sequenced the upstream regulatory region of *NaTPS38* in both Arizona and Utah genotypes. In comparison to the Utah genotype, the Arizona genotype has a 129 bp insertion and one single nucleotide polymorphism (SNP: change from A to G) within the 1 kb upstream region of *NaTPS38* (Fig. S1). In summary, polymorphisms at both *trans*- and *cis*-regulatory regions contribute to allelic differences of herbivory-induced *NaTPS38* expression and therefore (*E*)-α-bergamotene emission in *N. attenuata*.

**Fig. 5.**
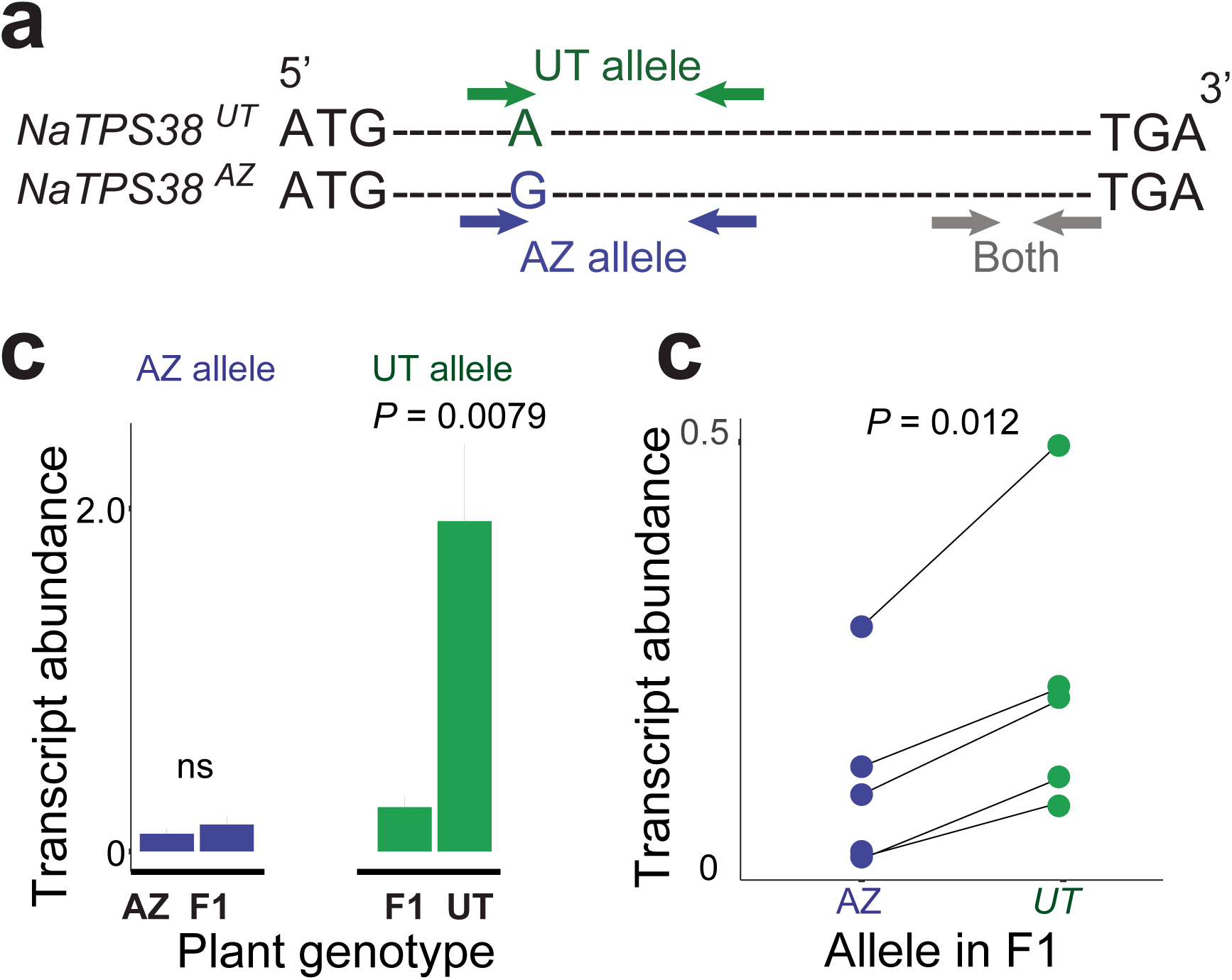
Allelic expression of *NaTPS38* in F1 hybrids of Arizona and Utah genotypes. (**a**) Scheme of primers used to measure the allele-specific transcript abundance. Green arrows refer to primers that can specifically amplify the Utah allele and blue arrows refer to primers that can specifically amplify the Arizona allele. Primers (grey arrows) that do not distinguish the two alleles were also used to measure total transcript abundance of *NaTPS38*. (**b**) Expression differences of *NaTPS38* alleles between F0 and F1 plants. Plants were treated with wounding plus oral secretion (N=5). Mean and SE are shown. Student’s *t*-test was used to determine differences between F0 and F1 plants. ns refers to no significant difference was found. (**c**) Expression differences of alleles in F1 individuals. Plants were treated with wounding plus oral secretion (N=5). Each dot represents an individual plant. Difference between two alleles was determined by paired Student’s *t*-test. Blue and green colors indicate alleles from Arizona and Utah genotypes, respectively.

### Allelic variations of *NaTPS25* and *NaTPS38* resulted in correlated changes of (*E*)-β-ocimene and (*E*)-α-bergamotene emission in HIPVs and floral volatiles in *N. attenuata*

We further explored whether the correlated changes in (*E*)-β-ocimene and (*E*)-α-bergamotene emission in HIPVs and floral volatiles are due to allelic variations of *NaTPS25* and *NaTPS38*. To this end, we first sequenced the full-length open reading frame of *NaTPS38* from the 25 natural genotypes. None of these contained either a premature stop codon or an insertion/deletion (Fig. 6**a**). The *NaTPS38* transcripts could be classified into three different haplotypes (YGI, HGV and YEI) that resulted from three common polymorphic non-synonymous substitution sites: Y32H, G280E, and I450V. Mapping these amino acid changes to the protein model of NaTPS38 suggested that Y32H and G280E are not involved in the formation of the active site cavity, though I450V is part of a loop structure near the entrance of the active site cavity (Fig. S2). However, the substitution I450V was found both in genotypes that emit high (e.g., genotype 304) and low levels (e.g., genotype 85) of herbivory-induced (*E*)-α-bergamotene, indicating that this amino acid change does not affect the enzymatic function of the NaTPS38 protein (Fig. 6**a**). We then compared (*E*)-α-bergamotene emission among natural genotypes that carry different haplotypes. The genotypes that carry the YEI-haplotype showed lower (*E*)-α-bergamotene emissions from both flowers and herbivory-induced leaves than those that harbour the YGI-haplotype (Fig. 6**b**). Consistently, abundance of *NaTPS38* transcripts from the YEI-haplotype carriers is also lower than those with the YGI-haplotype (Fig. S3). Interestingly, the HGV haplotype showed intermediate (*E*)-α-bergamotene emission in both flowers and herbivory-induced leaves (Fig. 6**b**). While the floral transcript abundance of *NaTPS38* in HGV-haplotype carriers is similar to that in YEI-haplotype carriers, in terms of herbivory-induced leaf abundances, they are similar to YGI-haplotype carriers (Fig. S3).

**Fig. 6.**
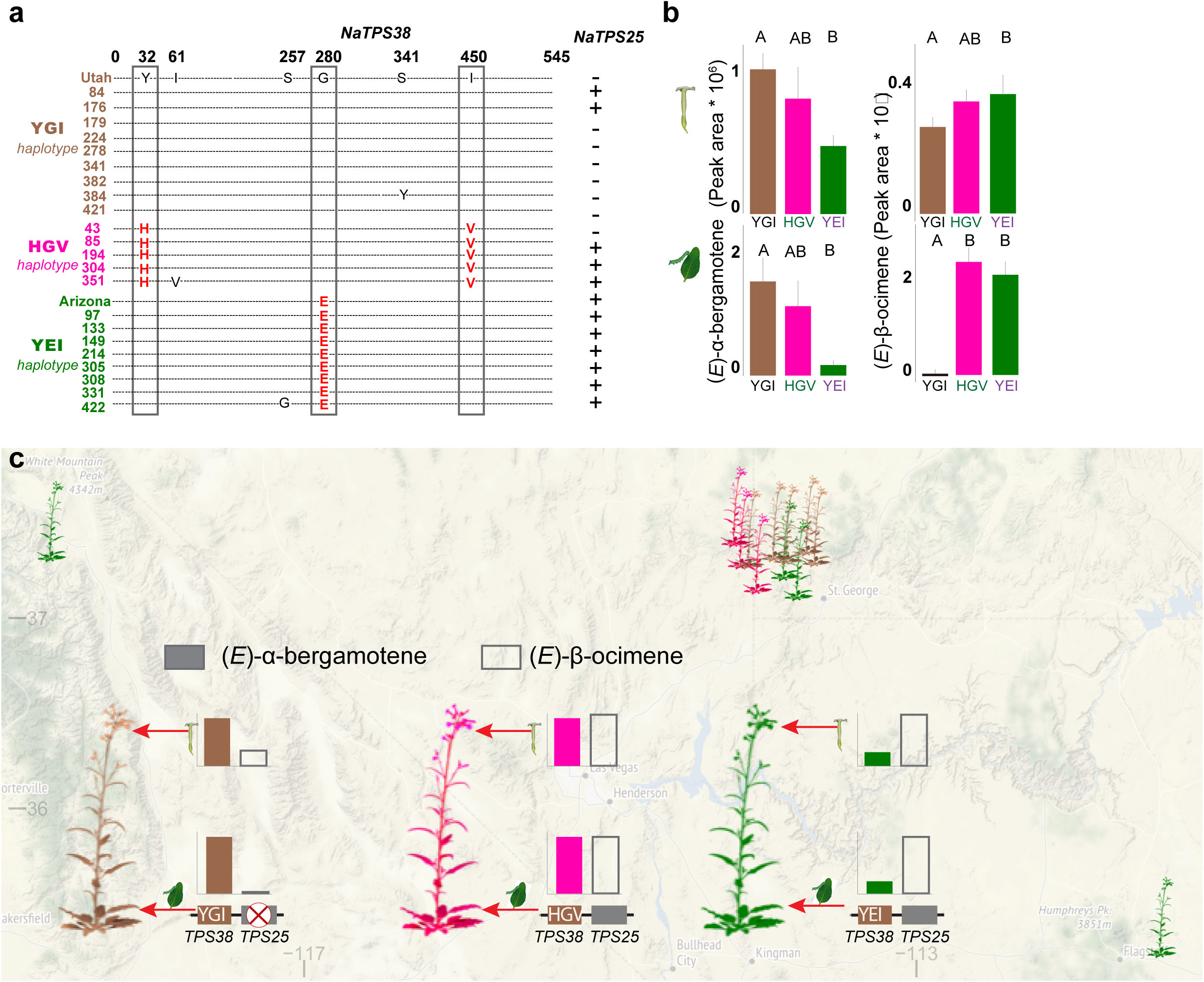
Allelic variations of the terpene cluster are associated with differences in floral volatiles and HIPVs among *N. attenuata* natural accessions. (**a**) Three different haplotypes found from the open reading frame of *NaTPS38* from *N. attenuata* accessions. The amino acid sequence of *NaTPS38* from the Utah genotype is set as the reference, and the different haplotype loci are marked in red. The presence of intact *NaTPS25* gene in the natural accessions are shown on the right (+, present; -not present). (**b**) The emission of (*E*)-α-bergamotene and (*E*)-β-ocimene in both flowers and herbivory-induced leaves are different among three *NaTPS38* haplotypes. The data for (*E*)-α-bergamotene emission were extract from our previous study (Zhou *et al*., 2017). Mean and SE are shown. Statistical differences among haplotypes are determined by ANOVA (*P* < 0.05). (**c**) A summary model of allelic differences of the two linked terpene synthases contribute to correlated intra-specific variation of floral and herbivory-induced volatiles in *Nicotiana attenuata*. The locations of different genotypes were shown on the map. Only a subset of individuals were shown near Utah. Colour represents different *NaTPS38* haplotype groups: brown refers to YGI, pink refers to HGV and green refers to YEI.

Because *NaTPS38* and *NaTPS25* are co-localized in the genome, we hypothesized that genetic variation of *NaTPS25* is also linked with the three different haplotypes of *NaTPS38*. Mapping *NaTPS25* presence/absence information in herbivory-induced leaves to the three haplotypes of *NaTPS38* showed that 80% (8 out of 10) of individuals that carried the *NaTPS38-YGI* haplotype also had no expression of *NaTPS25*, whereas individuals that carried the *NaTPS38-HGV* and *NaTPS38-YEI* haplotypes all had detectable *NaTPS25* transcript levels (Fig. 6**a**). Consistently, in both flowers and herbivory-induced leaves, individuals that carried the *NaTPS38-YGI* haplotype also accumulated only very low levels of *NaTPS25* transcripts and produced lower amount of (*E*)-β-ocimene, whereas individuals that carried *NaTPS38-YEI* haplotype accumulated high levels of *NaTPS25* transcripts and emitted high levels of (*E*)-β-ocimene (Fig. 3**a**, Fig. 6**b**, and Fig. S3**c**). Again, the *NaTPS38-HGV* haplotype plants showed intermediate *NaTPS25* transcript abundance but high levels of (*E*)-β-ocimene emission in both flowers and leaves (Fig. 3**a**, Fig. 6**b**, and Fig. S3**c**). These results reveal that the linked allelic differences of *NaTPS38* and *NaTPS25* in *N. attenuata* contributed to correlated changes in both floral volatiles and HIPVs.

## Discussion

Intrinsic genetic linkage and pleiotropy have been proposed to explain correlated variation in floral and defences traits (Johnson *et al*., 2015). However, direct genetic evidence for this is scarce. Here, by exploring natural variation and using both forward and reverse genetics tools, we demonstrate that allelic differences in two clustered terpene synthase genes, *NaTPS25* and *NaTPS38*, result in correlated intra-specific variations of both HIPVs and floral volatiles in *N. attenuata*.

We found that variations in the emissions of two terpenoids (*E*)-β-ocimene and (*E*)-α-bergamotene emissions showed correlated changes in HIPVs and floral volatile profiles (Figure 1). (*E*)-β-ocimene is often found among HIPVs and floral headspace in different species (Farre-Armengol *et al*., 2017). While (*E*)-β-ocimene in HIPVs can act as an important chemical cue for attracting natural enemies of phytophagous insects among plant species, it did not increase the attraction of predators of *M. sexta* larvae in *N. attenuata* in nature (Kessler & Baldwin, 2001). Floral (*E*)-β-ocimene emission is also common among flowering plants and can act as an attractant to generalist pollinators, such as bees and beetles etc. (Farre-Armengol *et al*., 2017). Although *N. attenuata* often attracts *M. sexta* moth as its pollinator, in some populations (e.g., Arizona), bees are also found to visit *N. attenuata* flowers (D. Kessler, personal communications). Therefore, it is plausible that floral (*E*)-β-ocimene emission in *N. attenuata* might increase bee-mediated pollination success. Different from (*E*)-β-ocimene, (*E*)-α-bergamotene emission is only reported in a few species, such as maize and *N. attenuata*. In both maize and *N. attenuata*, (*E*)-α-bergamotene in HIPVs is important for indirect defence function (Kessler & Baldwin, 2001; Degen *et al*., 2004). Although it remains unclear whether (*E*)-α-bergamotene is also emitted in floral tissues in maize, our resent study showed that the flowers of *N. attenuata* emit (*E*)-α-bergamotene, which increases *M. sexta* moths mediated pollination success (Zhou *et al*., 2017). Therefore, the correlated changes of (*E*)-β-ocimene and (*E*)-α-bergamotene in HIPVs and floral volatiles can result in concerted adaptation to pollinators and herbivores in *N. attenuata*.

The changes of (*E*)-β-ocimene and (*E*)-α-bergamotene emissions in HIPVs and floral headspace are likely due to allelic variations of two clustered terpene synthase genes, *NaTPS25* and *NaTPS38*. This is supported by four independent evidence. First, forward genetic mapping showed that the same locus is associated with the emissions of (*E*)-β-ocimene and (*E*)-α-bergamotene in HIPV. Although it was difficult to find the gene in the QTL directly from the fragmented *N. attenuata* genome assembly, synteny information between *N. attenuata* and tomato showed that the QTL likely contains a TPS cluster that includes *TPS25* and *TPS38*. Second, using reverse genetic approaches, we demonstrated that the expressions of *NaTPS25* and *NaTPS38* were required for the (*E*)-β-ocimene and (*E*)-α-bergamotene emission in HIPV (Fig. 2 and Zhou *et al*., 2017), respectively. Among different natural accessions, the expression levels of *NaTPS25* and *NaTPS38* were correlated with (*E*)-β-ocimene and (*E*)-α-bergamotene emission, respectively, in both floral headspace and HIPV (Fig. 2b & Zhou *et al*., 2017). Third, *in vitro* enzymatic assays also confirmed their specific terpene synthase functions (Fig. 2, Zhou *et al*., 2017). Fourth, allelic variations of *NaTPS38* and *NaTPS25* were tightly linked and formed three main haplotypes. Individuals carrying different haplotypes showed correlated changes of (*E*)-β-ocimene and (*E*)-α-bergamotene emissions in floral headspace and HIPV (Fig 6).

While the correlated changes of (*E*)-α-bergamotene in HIPVs and floral volatiles are due to pleiotropy of *NaTPS38*, the correlated changes of (*E*)-β-ocimene in HIPVs and floral volatiles are likely due to genetic linkage. This is because some genotypes do not emit (*E*)-β-ocimene in HIPVs due to loss-of-function mutations in *NaTPS25*, but they still emit (*E*)-β-ocimene in flowers, although at relatively low levels. A plausible scenario is that a transcriptional regulator of another TPS that is involved in floral (*E*)-β-ocimene emissions is in strong genetic linkage the *NaTPS25/38* locus. If this is the case, the correlation between emissions of (*E*)-β-ocimene in flower headspace and HIPV might disappear when the genetic linkage is interrupted, for example among genotypes that harbour much more frequent recombination than we have analysed.

We found two different genetic mechanisms are associated with allelic expression changes in *NaTPS25* and *NaTPS38*. While loss-of-function mutations mediated by a transposable element insertion resulted in expression variations of *NaTPS25*, changes at both *cis*- and *trans*-regulatory elements contributed to the different expressions of *NaTPS38* among genotypes. The differences in underlying genetic mechanisms might be due to differences in fitness optimal of (*E*)-α-bergamotene (one optimal peak, thus is under stabilizing selection) and (*E*)-β-ocimene (two optimal peaks, thus is under disruptive selection). However, it is currently unclear which stresses may favour loss-of-function in *NaTPS25*. Another explanation is that *NaTPS25* and *NaTPS38* have different pleiotropic functions. As silencing *NaTPS38* resulted in the complete absence of (*E*)-α-bergamotene in the HIPVs and floral volatile profiles (Zhou *et al*., 2017), NaTPS38 is likely the only key enzyme that is responsible for the biosynthesis of (*E*)-α-bergamotene in both HIPVs and floral volatiles in *N. attenuata*. However, while silencing *NaTPS25* resulted in strong decreases in (*E*)-β-ocimene emission in HIPVs, genotypes (such as Utah genotype) with loss-of-function in *NaTPS25* still emitted (*E*)-β-ocimene from flowers. This suggests that the biosynthesis of (*E*)-β-ocimene in flowers might be contributed by several genes with similar functions in *N. attenuata*. Therefore, the level of functional pleiotropy for *NaTPS25* is lower than for *NaTPS38*. As a consequence, a loss-of-function mutation in *NaTP25* might be associated with less distinct fitness costs than for one in *NaTPS38*. This is consistent with the pattern observed from the genetic mechanisms underlying changes of floral colour and scent (Xu *et al*., 2012; Wessinger & Rausher, 2012; Sheehan *et al*., 2016; Sas *et al*., 2016; Hoballah *et al*., 2007), in which loss-of-function mutations are more common than regulatory element changes in genes with less pleiotropy in their functions.

The intrinsic link between floral volatiles and HIPVs in natural populations of *N. attenuata* implies that intra-specific variation of plant volatiles can be maintained due to pleiotropy and/or genetic linkage. For example, when a *N. attenuata* population is under directional selection for high amounts of floral (*E*)-α-bergamotene emission, either imposed by *M. sexta* moths or hummingbirds (both show strong preference for (*E*)-α-bergamotene (see Kessler & Baldwin, 2007)), individuals that carry the YGI haplotype would have higher fitness and the frequency of the YGI haplotype in the population would increase. As a consequence, the level of herbivory-induced (*E*)-α-bergamotene and (*E*)-β-ocimene emissions would increase in the population. In such a scenario, changes in HIPVs in these population are not due to selection on HIPVs, but rather owing to the pleiotropy or genetic linkage of the YGI haplotype, which is also associated with high levels of herbivory-induced (*E*)-α-bergamotene and low levels of (*E*)-β-ocimene emission. The same logic may also apply to how selection on HIPVs could result in changes in floral volatiles.

The presence of genetic linkage and pleiotropy between HIPVs and floral volatiles suggest the evolution of plant volatiles might evolve under diffuse selection imposed from herbivores and pollinators in *N. attenuata* (Iwao & Rausher, 1997; Strauss & Irwin, 2004; Agrawal & Stinchcombe, 2009; Wise & Rausher, 2013). Diffuse coevolution can be common for plant resistance to multiple-herbivore communities (Wise & Rausher, 2013) and recent genetic analysis have discovered correlated changes of chemical defence compounds among tissues (Chan *et al*., 2011; Keith & Mitchell-Olds, 2019) were due to genetic linkage or pleiotropy, similar to the two terpenoids we found here. Although future studies that use comparative metabolomics and transcriptomics are needed to further examine this, it is plausible that many other metabolic traits also show correlated changes among tissues. The diffuse interactions to multiple-herbivore communities and correlated changes of chemical defences among tissues often constrain the evolution of plant resistance (Wise & Rausher, 2013; Keith & Mitchell-Olds, 2019). Because plant-herbivore and plant-pollinator interactions were often studied in isolation (Lucas-Barbosa, 2016), to extent which diffused interaction exists among plant-herbivore and plant pollinator interactions, and what are the evolutionary consequences remain largely unclear. However, a recent study using an experimental evolution approach has demonstrated a diffuse coevolution of floral signals and plant direct defences in *Brassica rapa* (Ramos & Schiestl, 2019), although this study did not investigate the levels of genetic correlation of floral signals and plant defences. These studies and results reported here highlighted the importance of investigating traits evolution in a community context (Strauss & Irwin, 2004). Prospective studies deploying experimental evolution approaches, in which the multi-generational fitness of different genotypes is determined under manipulated selection agents in their native habitat, will be needed to provide complementary understanding on how adaptive traits evolve in nature.

## Abbreviations

HIPVs: herbivory-induced plant volatiles
TPS: terpene synthase
VIGS: virus induced gene silencing
AI-RIL: advanced inter-cross recombinant inbred line

## Acknowledgements

We thank Dr. M. Huber and Dr. M. Schäfer for critical comments on an earlier version of the manuscript. We are grateful to the help from Anke Kügler for sample collections. This work was supported by funding from the Swiss National Science Foundation (PEBZP3-142886 to S.X.), a European Commission Marie Curie Intra-European Fellowship (IEF, 328935 to S.X.), the Max Planck Society, and a European Research Council Advanced Grant (ClockworkGreen 293926 to I.T.B.).

## Authors’ contributions

Conceptualization, S.X.; Methodology, W.Z., S.X., C.K., E.M., N.D.L., H.G., T.G.K., I.T.B. and M.C.S; Investigation, W.Z., S.X., C.K., E.M., N.D.L., H.G., T.G.K., and M.C.S.; Writing, S.X., W.Z.; Review and Editing, S.X., W.Z., C.K., E.M., N.D.L., H.G., T.G.K., M.C.S., and I.T.B.; Funding Acquisition, S.X., I.T.B.; Resources, S.X. and I.T.B.

## Supplementary Material

### Supplementary Figures

**Fig. S1 The differences in the putative promoter region of *NaTPS38* between Arizona and Utah genotypes**. The 5’ upstream genomic sequences of *NaTPS38* are shown. The putative TATA box is bounded by a red box. Presumed tandem repeats are enclosed within the blue box.

**Fig. S2 Protein Structure model of NaTPS38**. The homology-based model was created using the crystal structure of 1,8-cineol synthase from *Salvia fruticosa* (PDB 2j5c.1) as template using the web-based swiss model service (https://swissmodel.expasy.org). Tyrosine 32, glycine 280, and isoleucine 450 are shown in brown. The active site cavity is marked in yellow.

**Fig. S3 Transcript abundance of *NaTPS38* and *NaTPS25* alleles among different NaTPS38 haplotypes in *N. attenuata***. (**a**) The relative transcript abundance of *NaTPS38* in the flowers of different *NaTPS38* haplotypes. (**b**) The relative transcript abundance of *NaTPS38* in the herbivory-induced leaves of different *NaTPS38* haplotypes. (**c**) The relative transcript abundance of *NaTPS25* in the flowers of different *NaTPS38* haplotypes. Mean and SE are shown. Statistical differences among haplotypes are determined by ANOVA (*P* < 0.05).

### Supplementary Tables

**Table S1.** GPS coordinates of location where the 25 *N. attenuata* natural accessions were originally collected.

| Accession | Latitude | Longitude |
| --- | --- | --- |
| 43 | 37°17'09.10"N | 114°7'31.50"W |
| 84 | 37°19'35.48"N | 113°57'38.28"W |
| 85 | 37°19'35.48"N | 113°57'38.28"W |
| 97 | 37°21'35.24"N | 113°56'38.68"W |
| 133 | 37°06'12.50"N | 113°49'36.60"W |
| 138 | 37°08'26.36"N | 114°01'39.41"W |
| 149 | 35°12'56.07"N | 111°27'41.29"W |
| 176 | 37°16'38.65"N | 113°53'35.18"W |
| 179 | 37°21'01.04"N | 113°57'05.17"W |
| 194 | 37°20'22.52"N | 114°02'40.86"W |
| 214 | 37°13'15.83"N | 113°48'20.86"W |
| 224 | 37°19'33.89"N | 113°57'54.16"W |
| 278 | 37°16'16.22"N | 114°07'39.67"W |
| 304 | 37°20'22.52"N | 114°02'40.86"W |
| 305 | 37°45'19.61"N | 118°35'41.82"W |
| 308 | 37°13'05.50"N | 113°48'24.25"W |
| 331 | 37°13'15.83"N | 113°48'20.86"W |
| 341 | 37°09'45.30"N | 114°00'58.52"W |
| 351 | 37°17'09.10"N | 114°07'31.50"W |
| 382 | 37°14'27.05"N | 113°49'36.71"W |
| 384 | 37°14'27.05"N | 113°49'36.71"W |
| 421 | 37°07'11.90"N | 114°00'25.10"W |
| 422 | 35°12'44.80"N | 111°28'24.80"W |
| Utah<br>(inbred) | 37°19'36.26"N | 113°57'53.05"W |
| Arizona<br>(inbred) | 35°13'08.62"N | 111°28'26.03"W |

**Table S2.**
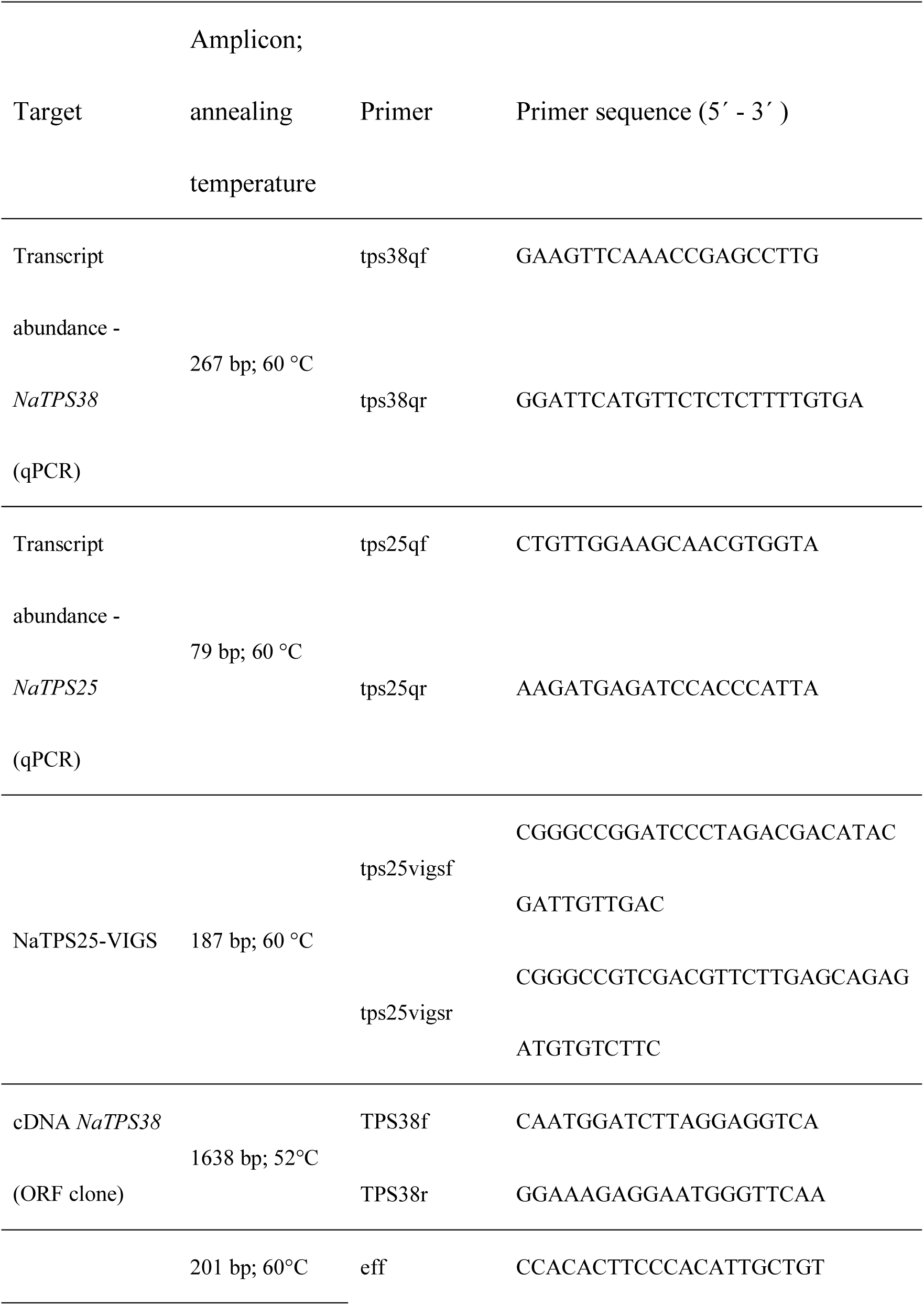
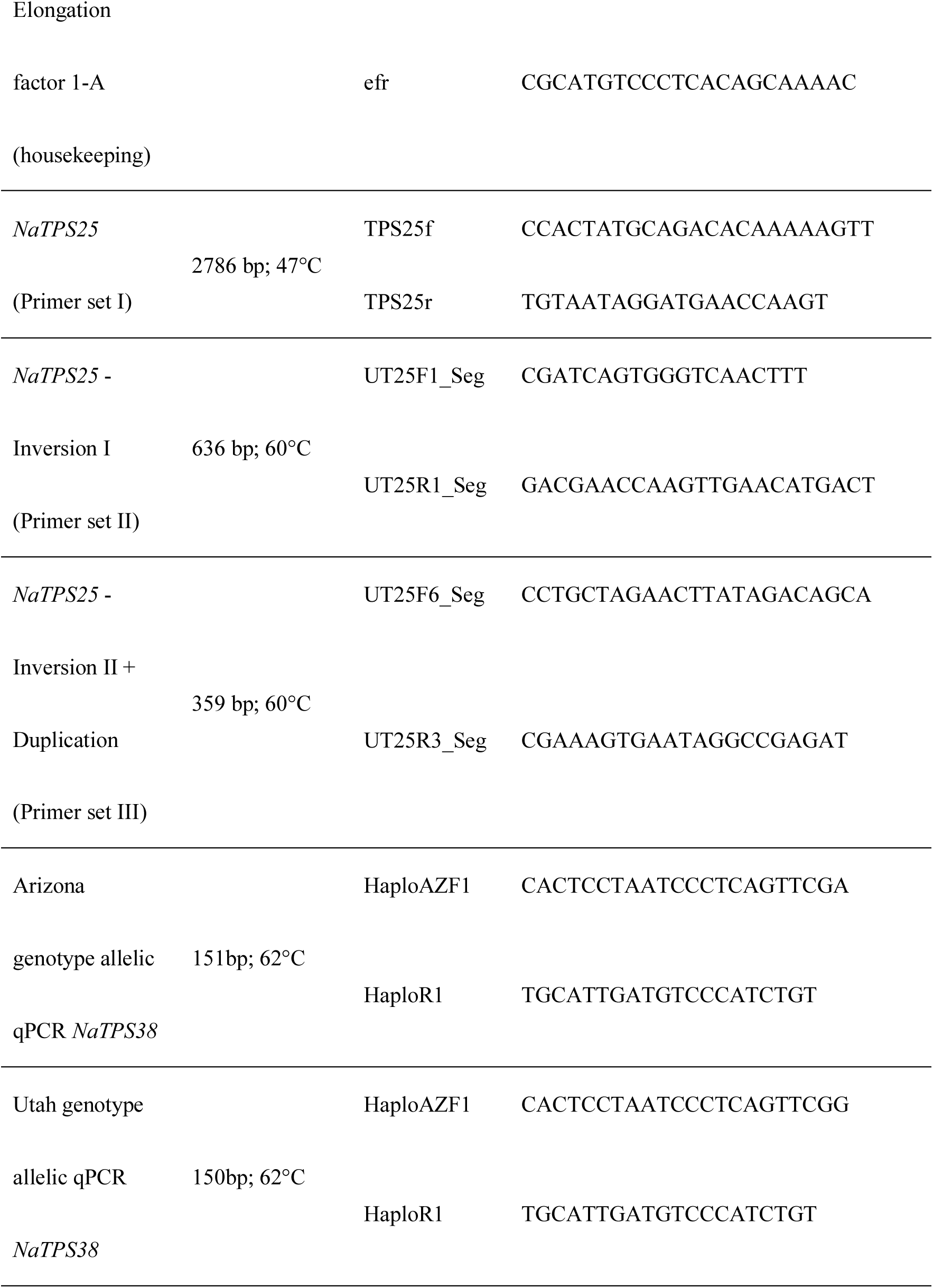
The primer sets used in this study.

