## Supplementary figures and images for "Allelic differences of clustered terpene synthases contribute to correlated intra-specific variation of floral and herbivory-induced volatiles in a wild tobacco"

### Fig. S1

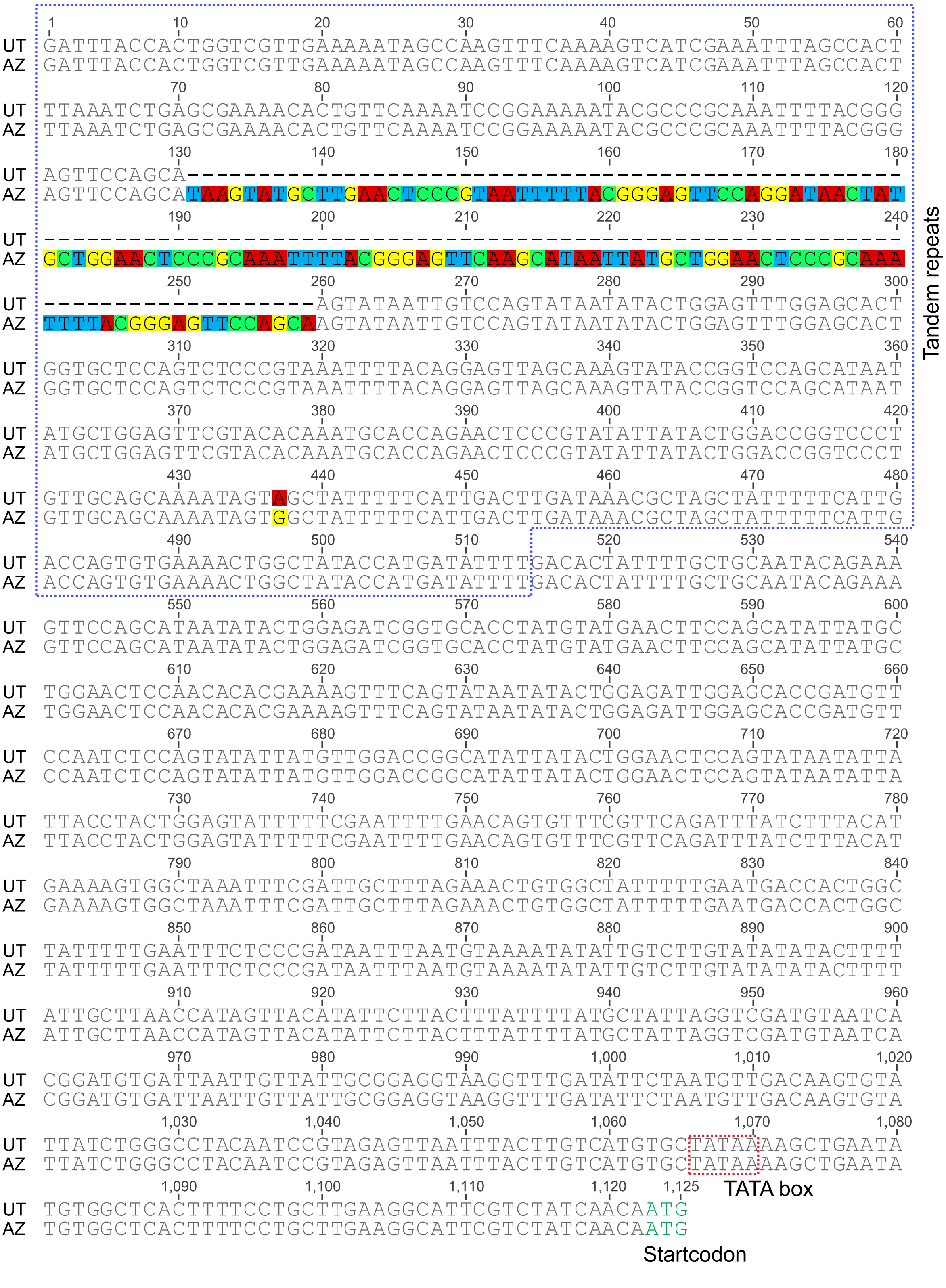

### Fig. S2

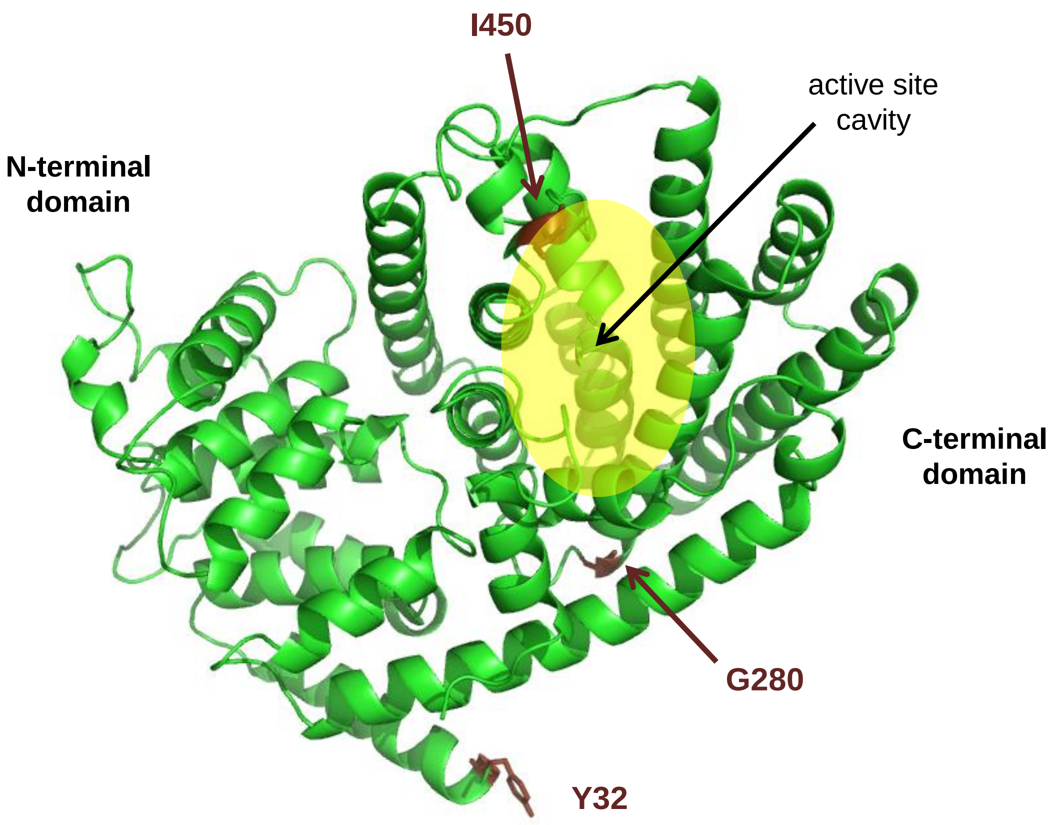

### Fig. S3

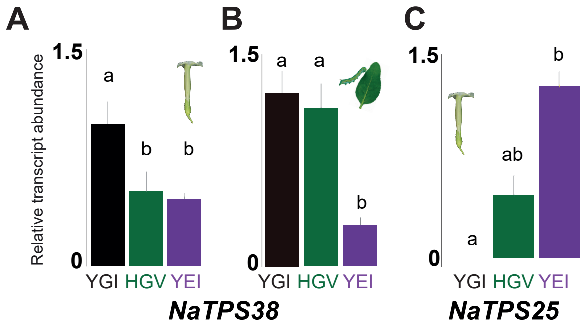
